# PSAP-genomic-regions: a method leveraging population data to prioritize coding and non-coding variants in whole genome sequencing for rare disease diagnosis

**DOI:** 10.1101/2024.02.13.580050

**Authors:** Marie-Sophie C. Ogloblinsky, Ozvan Bocher, Chaker Aloui, Anne-Louise Leutenegger, Ozan Ozisik, Anaïs Baudot, Elisabeth Tournier-Lasserve, Helen Castillo-Madeen, Daniel Lewinsohn, Donald F. Conrad, Emmanuelle Génin, Gaëlle Marenne

## Abstract

The introduction of next generation sequencing technologies in the clinics has improved rare disease diagnosis. Nonetheless, for very heterogeneous or very rare diseases, more than half of cases still lack molecular diagnosis. Novel strategies are needed to prioritize variants within a single individual. The PSAP (Population Sampling Probability) method was developed to meet this aim but only for coding variants in exome data. To address the challenge of the analysis of non-coding variants in whole genome sequencing data, we propose an extension of the PSAP method to the non-coding genome called PSAP-genomic-regions. In this extension, instead of considering genes as testing units (PSAP-genes strategy), we use genomic regions defined over the whole genome that pinpoint potential functional constraints.

We conceived an evaluation protocol for our method using artificially-generated disease exomes and genomes, by inserting coding and non-coding pathogenic ClinVar variants in large datasets of exomes and genomes from the general population.

We found that PSAP-genomic-regions significantly improves the ranking of these variants compared to using a pathogenicity score alone. Using PSAP-genomic-regions, more than fifty percent of non-coding ClinVar variants, especially those involved in splicing, were among the top 10 variants of the genome. In addition, our approach gave similar results compared to PSAP-genes regarding the scoring of coding variants. On real sequencing data from 6 patients with Cerebral Small Vessel Disease and 9 patients with male infertility, all causal variants were ranked in the top 100 variants with PSAP-genomic-regions.

By revisiting the testing units used in the PSAP method to include non-coding variants, we have developed PSAP-genomic-regions, an efficient whole-genome prioritization tool which offers promising results for the diagnosis of unresolved rare diseases. PSAP-genomic-regions is implemented as a user-friendly Snakemake workflow, accessible to both researchers and clinicians which can easily integrate up-to-date annotation from large databases.

**Author summary:** In recent years, improvement in DNA sequencing technologies has allowed the identification of many genes involved in rare diseases. Nonetheless, the molecular diagnosis is still unknown for more than half of rare diseases cases. This is in part due to the large heterogeneity of molecular causes in rare diseases. This also highlights the need for the development of new methods to prioritize pathogenic variants from DNA sequencing data at the scale of the whole genome and not only coding regions. With PSAP-genomic-regions, we offer a strategy to prioritize coding and non-coding variants in whole-genome data from a single individual in need of a diagnosis. The PSAP-genomic-regions combines information on the predicted pathogenicity and frequency of variants in the context of functional regions of the genome. In this work, we compare the PSAP-genomic-regions strategy to other variant prioritization strategies on simulated and real data. We show the better performance of PSAP-genomic-regions over a classical approach based on variant pathogenicity scores alone. PSAP-genomic-regions provides a straightforward approach to prioritize causal pathogenic variants, especially non-coding ones, that are often missed with other strategies and could explain the cause of undiagnosed rare diseases.

## Introduction

Each rare disease affects, by definition, a small number of individuals. However, as a whole, rare diseases affect about 350 million people world-wide (1). Approximately 80% of rare diseases have a genetic origin that mostly follows a Mendelian mode of inheritance (2–4). The advent of Next Generation Sequencing (NGS) and the development of variant pathogenicity prediction tools have allowed, in recent years, the identification of many genes involved in rare Mendelian diseases. Nonetheless, despite extensive efforts, the molecular diagnosis is still unknown for more than 50% of rare diseases cases (5–7). This can mainly be explained by the fact that many rare diseases are characterized by an extreme genetic heterogeneity, which results in only one individual carrying a specific pathogenic causal variant. This issue is referred to as the “n-of-one” problem (8).

With the advent of high throughput sequencing technologies in clinics, molecular diagnosis is now often sought through whole exome or whole genome sequencing (WES and WGS respectively). However, due to the large number of rare variants in each individual genome, causal variants are sought among very rare and highly pathogenic variants in genes relevant to the current known disease mechanism. The limited knowledge about gene functions and disease mechanisms can make this strategy unfruitful. To address the issue of variant prioritization at the level of an individual, the Population Sampling Method (PSAP) (8) was developed. PSAP computes, for each gene, a null distribution, which is the probability to observe in the general population a genotype with a CADD pathogenicity score (9) greater than or equal to the highest one to the highest one observed in the patient for this gene. This initial version of the PSAP method, which we will refer to as PSAP-genes, has been successfully applied to identify variants of interest in diverse phenotypes, including male infertility (10–12), recurrent pregnancy loss (13) and ciliary diskynesia (14).

A current hindrance to the application and generalization of PSAP-genes as a tool for diagnosis is its restriction to the coding parts of the genome. Indeed, the majority of variants reside in non-coding parts of the genome (15). Non-coding variants may contribute to explain part of the etiology of rare diseases (16), as suggested by the large number of GWAS hits located in non-coding regions of the genome (17). The involvement of non-coding pathogenic variants in rare diseases is further corroborated by the fact that non-coding regions are heavily involved in the regulation of gene expression. Several prediction tools have been developed to this end (18–20), but most of them lack a variant-based score for both coding and non-coding regions. In addition, to be performant, they often require multiple annotations like Human Phenotype Ontology (HPO) terms (21) to characterize the symptoms or disease of a patient . Thus, they rely on previous knowledge and rarely go beyond candidate genes.

To move beyond the gene as a natural unit of testing for the PSAP method, we need to use predetermined regions across the whole genome. These regions also need to be defined using functional information to be used as a cohesive unit for the construction of PSAP null distributions. This challenge of defining regions along the whole genome has been tackled by Bocher et al. in the context of rare-variant association testing (22): they describe CADD regions, which are characterized by a lack of observed variants with high functionally-Adjusted CADD Scores (ACS) in the gnomAD database (23). CADD regions are expected to reflect functional constraints. CADD regions present the key advantage of providing pre-defined and functionally-informed regions which can be used to construct PSAP null distributions.

We have made available a new implementation of the PSAP method using Snakemake (24) workflows, called Easy-PSAP (https://github.com/msogloblinsky/Easy-PSAP), which features null distributions constructed with up-to-date allele frequency data and pathogenicity scores. Here, we introduce PSAP-genomic-regions, an extension of the PSAP method to the non-coding genome by using the pre-defined CADD regions as testing unit instead of genes. This is an innovative strategy to prioritize variants at the scale of an individual genome. PSAP-genomic-regions is now available in Easy-PSAP. We devised an evaluation protocol using artificially-generated disease exomes and genomes, obtained by inserting coding and non-coding ClinVar (25) variants in general population whole genomes from the 1000 Genomes Project (26) and exomes from the FrEnch EXome (FREX) project (27). We show the consistent improvement in prioritization by using PSAP-genomic-regions over pathogenicity scores alone for non-coding and then coding variants. For coding variants, we also demonstrate the good performance of PSAP-genomic-regions compared to PSAP-genes. On real-life data, we illustrate the power of PSAP-genomic-regions on WES data from six resolved cases of Cerebral Small Vessel Disease (CSVD) and WGS data from three families affected by male infertility. These two diseases are particularly relevant to test our method, monogenic forms of CSVD (28) and male infertility (29) being extremely heterogeneous.

## Results

### Construction of PSAP null distribution in coding and non-coding regions

The idea behind the original PSAP method, referred to as PSAP-genes, relies on the calculation of gene-specific null distributions of CADD pathogenicity scores. More precisely, for an individual exome or genome and in a given gene, PSAP-genes considers the genotype with the highest CADD score and evaluates the probability to observe such a high CADD score in this gene in the general population (see S1 File for a detailed explanation of the calculation of PSAP null distributions). PSAP-genes deals separately with heterozygote and homozygote variants in the autosomal dominant (AD) and the autosomal recessive (AR) models respectively. As a result, PSAP-genes gives a p-value to the genotype with the highest CADD score in the gene for each gene, model, and individual. This p-value allows the ranking of the genes for an individual exome or genome. The PSAP principle can be generalized to any genomic unit.

Here, with PSAP-genomic-regions, we extended the PSAP method to analyze whole-genome data using predefined CADD regions as testing units instead of genes (Fig 1). The same principle as before is employed, with the difference being that the genotype with the highest CADD score in the region can be coding or non-coding. We thus constructed PSAP-genomic-regions null distributions with two pathogenicity scores : the initial CADD score (PHRED scaled across the whole genome), or the ACS (22) (PHRED scaled CADD scores by “coding”, “regulatory” and “intergenic” regions) to mitigate the higher CADD scores of coding variants. Our two novel strategies will be referred to as PSAP-genomic-regions-CADD and PSAP-genomic-regions-ACS. They were compared to the initial PSAP-genes strategy, also referred to as PSAP-genes-CADD.

**Fig 1.**
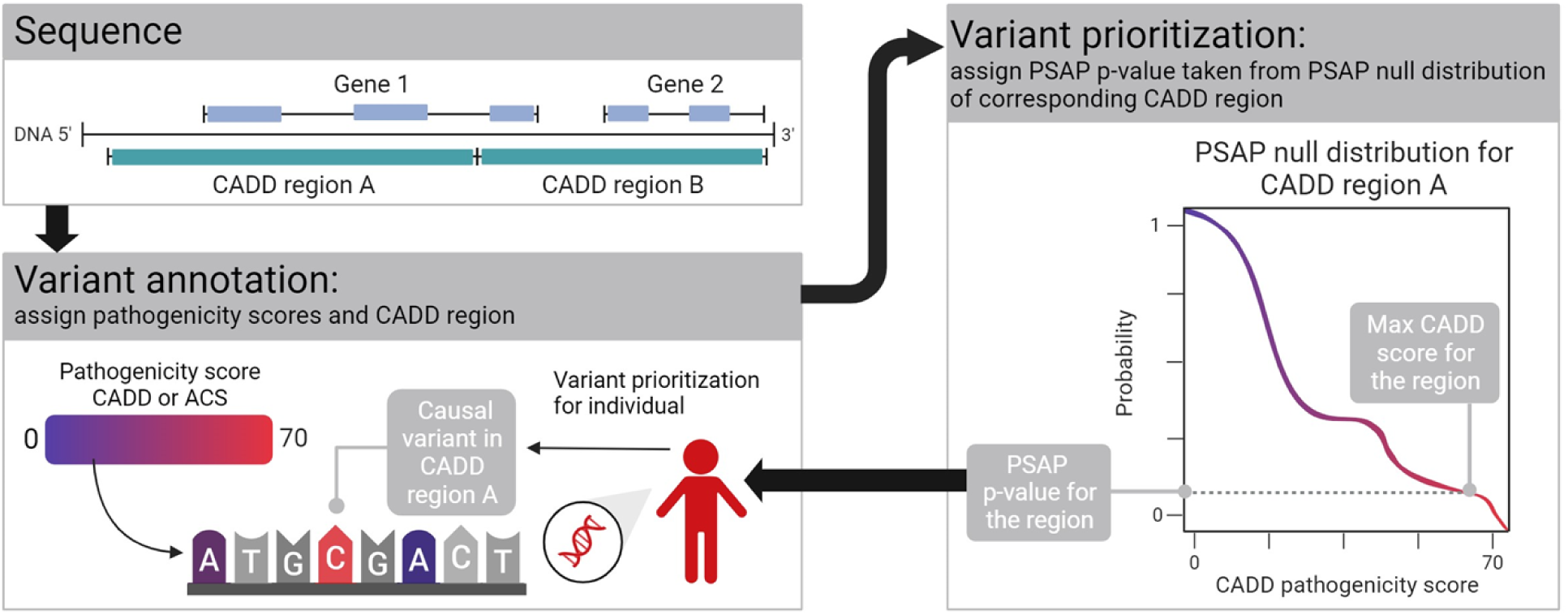
Description of the PSAP-genomic-regions strategy. We calculated PSAP null distributions for SNVs in genes and CADD regions, in the hg19 and hg38 assemblies of the human genome. In hg19, PSAP null distributions were obtained for 19,283 genes and 119,695 CADD regions. In hg38 PSAP null distributions were obtained for 18,395 genes and 123,991 CADD regions. PSAP null distributions and their parameters (unit of testing, allele frequencies and pathogenicity score) can be found in S1 Table.

### Evaluating the performance of PSAP-genomic-regions on artificially-generated disease exomes and genomes using ClinVar variants

#### Prioritization of non-coding pathogenic variants

First, to evaluate how PSAP-genomic-regions performed to prioritize non-coding pathogenic variants, we used artificially-generated disease genomes created by inserting non-coding ClinVar variants in the NFE genomes from 1000G project (see Material & Methods and S2 File for the list of variants). Because the 1000 Genomes project is population-based, we expect that some individuals might carry one or a few pathogenic variants in their genome. These pathogenic variants are characterized by a high CADD score and a low PSAP p-value. Indeed, there is large variation in the maximal CADD score or lowest PSAP p-value, whereas the rest of the distribution is extremely similar between individuals (S1 Fig). Thus, in order to summarize the rank of a ClinVar variant in an evaluation setting, we considered the best rank reached by the variant in at least 90% of the individuals.

Most of the NFE genomes carried a variant with a higher pathogenicity score or a lower PSAP p-value than most of the ClinVar variants (S2 Fig). We thus compared the percentage of the non-coding pathogenic variants ranked among the top N (N = 1, 10, 50 and 100) in at least 90% of the NFE genomes. The ranking at the individual level was done among all heterozygous variants for the ClinVar variants under the AD model, and across homozygous variants for the ClinVar variants under the AR model. (Fig 2A). With both CADD and ACS pathogenicity scores, PSAP-genomic-regions performed systematically better than using the pathogenicity scores alone. The improvement was especially large for the top 10 ranking: 24.6% and 79.2% of ClinVar variants reached the top 10 with PSAP-genomic-regions-CADD for the AD and AR models, respectively, while no ClinVar variant reached the top 10 with CADD scores alone.

**Fig 2.**
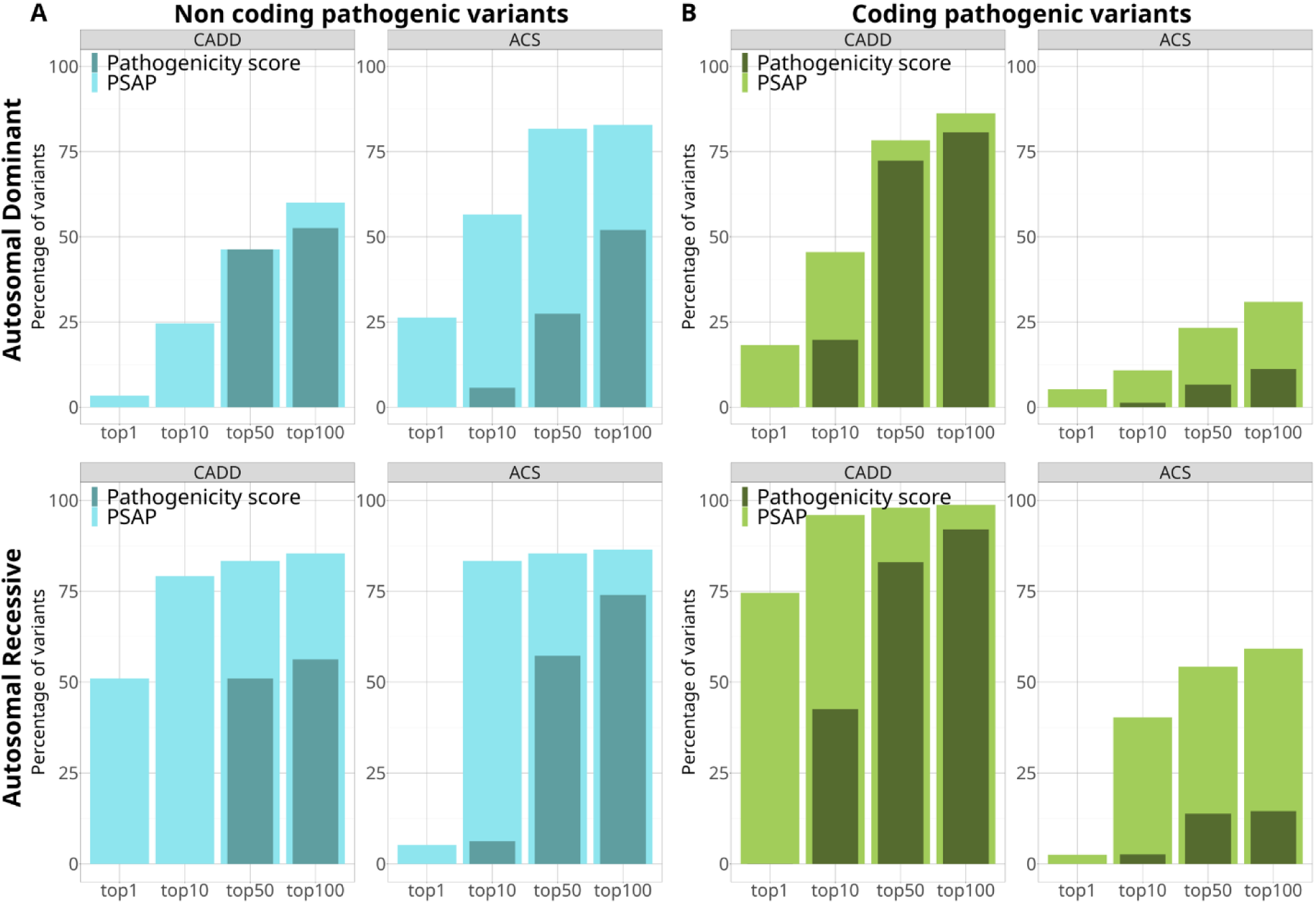
Comparison of the PSAP-genomic-regions strategy versus a pathogenicity score alone for in artificially-simulated disease genomes. Percentage of non-coding and coding pathogenic ClinVar variants reaching the top N of variants in at least 90% of NFE genomes, with PSAP-genomic-regions (darker shade of blue or green) or the pathogenicity score alone (lighter shade of blue or green), CADD or ACS (A) N = 175 non-coding AD variants and N = 96 non-coding AR variants (B) N = 4,965 coding AD variants and N = 2,680 coding AR variants.

Using the ACS scores further improved the performance to detect non-coding-variants: 56.6% and 83.3% of variants reached the top 10 with PSAP-genomic-regions-ACS for the AD and AR models, respectively. Nonetheless, we can note the pattern is different for the top 1 for the AR model: 51% with PSAP-genomic-regions-CADD to 5.5% with PSAP-genomic-regions-ACS. Indeed, switching from CADD score to ACS score has lowered the PSAP p-value of non-coding variants shared by more than 10% of NFE genomes. This led to a defect of the top rank reached by the ClinVar variants, as we considered the lowest rank reached in at least 90% of individuals. For instance, a variant in the CADD region R109138 shared by 70 of the NFE genomes went from a CADD score of 18.1 and a PSAP-genomic-regions-CADD p-value of 0.1 to an ACS of 22.2 and a PSAP-genomic-regions-ACS p-value of 5.18x10^-10^. Thus, the ClinVar variants inserted in these individuals having a higher p-value than 5.18x10^-10^ do not rank first.

We further explored PSAP results for splicing ClinVar variants versus other type of non-coding ClinVar variants. Indeed, we observed that splicing variants are the major type of non-coding ClinVar variants. These splicing variants often had a very good ranking, especially with PSAP-genomic-regions-ACS (n=115 splicing variants among 175 non-coding AD variants and n=72 splicing variants among 96 non-coding AR variants; S3 Table; Panel A in S3 Fig). Splicing ClinVar variants have a much higher ACS than CADD scores (Panel B in S3 Fig) which results in better ranking than for other types of non-coding ClinVar variants using PSAP-genomic-regions-ACS p-values (Panel C in S3 Fig). As a consequence, the percentage of splicing ClinVar variants ranked in the top 10 was largely improved when using PSAP-genomic-regions-ACS, for the AD model especially which was less powerful with PSAP-genomic-regions-CADD to begin with (Panel D in S3 Fig).

The full results of ranking by PSAP-genomic-regions-ACS for the non-coding non-splicing pathogenic ClinVar variants can be found in S3 File. With PSAP-genomic-regions-ACS, around half of the non-coding non-splicing variants are ranked in the top 100 of variants for more than 90% of NFE genomes (46 out of 73 variants for the AD model and 19 out of 31 variants for the AR model). The other half of variants present a less significant PSAP-genomic-regions-ACS p-value and a poorer ranking. To confirm this pattern of ranking for non-coding non-splicing pathogenic variants on another set of variants, we evaluated with our artificially generated disease genomes protocol 320 non-coding SNVs used to train Genomiser (30). These variants were not associated with a mode of inheritance. Hence, we inserted them in the NFE genomes and scored them with both AD and AR PSAP-genomic-regions-ACS null distributions. Among the 320 non-coding variants, 169 reached the top 100 in at least 90% of NFE genomes, with either the AD or AR model (S4 File). This can be explained by the distributions of CADD scores compared to ACS scores for the ClinVar variants: the non-coding variants that do not reach the top 100 have a significantly lower CADD and ACS scores compared to all the other types of variants (S4 Fig). Overall, PSAP-genomic-regions-ACS prioritizes around half of non-coding ClinVar and Genomiser training variants in the top 100 of NFE genomes. The ones who have a higher ranking present much lower CADD and ACS scores and would never be well-ranked by any PSAP strategy.

PSAP-genomic-region is also relevant for the analysis of exome data. Indeed, exome sequencing captures variants outside of the bounds of coding regions (31), such as intronic variants. We explored the prioritization of non-coding ClinVar variants located within the WES-targeted regions of the FREX individuals using our artificially-generated disease exomes protocol (N=48 variants for the AD model and N=64 variants for the AR model, Panel A in S5 Fig). For both PSAP-genomic-regions-CADD and PSAP-genomic-regions-ACS, there was a large increase in prioritization performance compared to using only the pathogenicity scores. Because there are fewer variants in an exome background than in a genome background, the rankings of these non-coding ClinVar variants were better in FREX than in NFE genomes. The best ranking was achieved using PSAP-genomic-regions-ACS, with 82% and 90.3% of variants reaching the top 10 for the AD and AR models, respectively. Most of these non-coding pathogenic variants were splicing variants (40 out of 73 variants for the AD model and 56 out of 64 variants for the AR model), and half of them were considered as having a functional “HIGH IMPACT” (26 variants for the AD model and 22 variants for the AR model). Hence, prioritizing variants with PSAP on CADD regions allows identifying more variants even in exome data, that are in addition functionally-relevant.

#### Prioritization of coding pathogenic variants

Similar evaluations were performed for ClinVar coding variants inserted in either WGS from 1000G NFE individuals or WES from FREX. As observed for non-coding pathogenic variants, PSAP-genomic-regions outperformed the pathogenicity scores alone (Fig 2B, Panel B in S5 Fig). However, in the context of coding pathogenic ClinVar variants, we observed that the strategy of PSAP-genomic-regions-CADD provided better prioritization compared with the PSAP-genomic-regions-ACS strategy. We observed that 18.2% and 74.6% of the coding variants reached the top 1 in at least 90% of genomes backgrounds with the PSAP-genomic-regions-CADD for the AD and AR model respectively, against no variants with the CADD score alone, and against 5.3% and 2.5% reaching the top 1 with PSAP-genomic-regions-ACS. In the exome background and with PSAP-genomic-regions-CADD, 38.7% and 89.8% of AD variants reached the top 1 and top 50, respectively; 80.3% and 97.9% of AR variants reached the top 1 and the top 50, respectively.

We also compared the number of coding ClinVar variants reaching the tops in NFE genomes between PSAP-genomic-regions-CADD strategy and the initial PSAP-genes-CADD strategy (Fig 3). More differences were observed across the two PSAP strategies for the AD than for the AR model (Fig 3A). There were 362 variants ranked first and 1,017 variants ranked [2-10] in common between the two strategies. However, 908 variants that were ranked [2-10] with PSAP-genes-CADD were [11-50] with PSAP-genomic-regions-CADD, and 395 variants that were ranked [2-10] with PSAP-genes-CADD were ranked first with PSAP-genomic-regions-CADD. Regarding variants that are ranked more than a 100 with PSAP-genomic-regions-CADD, 278 of them are ranked [11-50] and are ranked [51-100] by PSAP-genes-CADD. Regarding the AR model (Fig 3B), PSAP-genomic-regions-CADD performed similarly to PSAP-genes-CADD, and the majority of variants were ranked first with both strategies (1,550 variants). Even more promising results can be found when looking at the same comparison of ranks within the FREX exomes (S6 Fig). For instance, in the AD model, 592 variants that were ranked [2-10] with PSAP-genes-CADD are ranked first with PSAP-genomic-regions-CADD, against 115 variants ranked [2-10] with PSAP-genomic-regions-CADD that become first with PSAP-genes-CADD.

**Fig.3.**
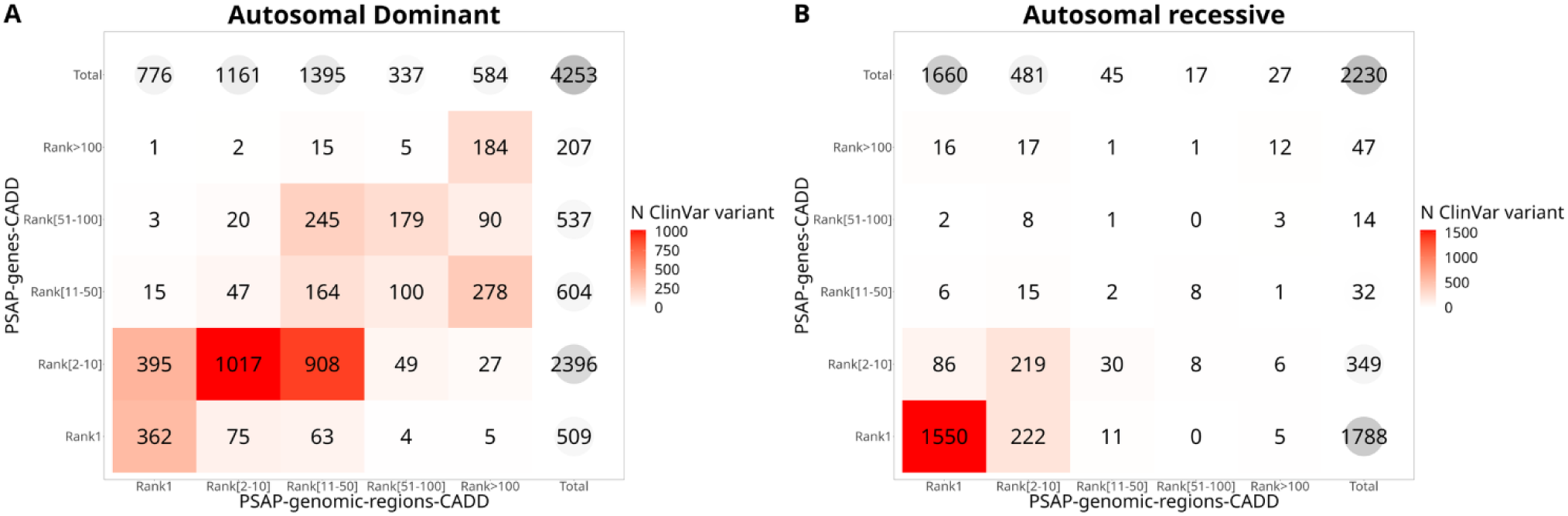
Comparison of PSAP-genomic-regions-CADD and PSAP-genes-CADD strategies in artificially-simulated disease genomes. Number of coding pathogenic ClinVar variants reaching rank [x-y] of variants in at least 90% of 1000 Genomes Project NFE individuals for each strategy.

### Application of PSAP-genomic-regions to real data with different modes of inheritance

To illustrate our method in real-life settings, we analyzed two datasets (S4 Table), one with an AD mode of inheritance and the other with an AR mode of inheritance. The first dataset consisted of WES data for six individuals affected by monogenic forms of CSVD (32). Using PSAP-genomic-regions-CADD, all of the causal variants were ranked at least in the top 100 in each patient (Fig 4). The contribution of CADD regions as a unit of testing was especially visible for the variant in *COL4A2* and one variant in *HTRA1* which were not well-ranked using genes as testing unit (rank 110 and 193 respectively with genes, and rank 3 and 69 with CADD regions). Using their maximal CADD score by gene or CADD region alone, these variants would not have been prioritized in the top 100 for five out of six individuals.

**Fig. 4.**
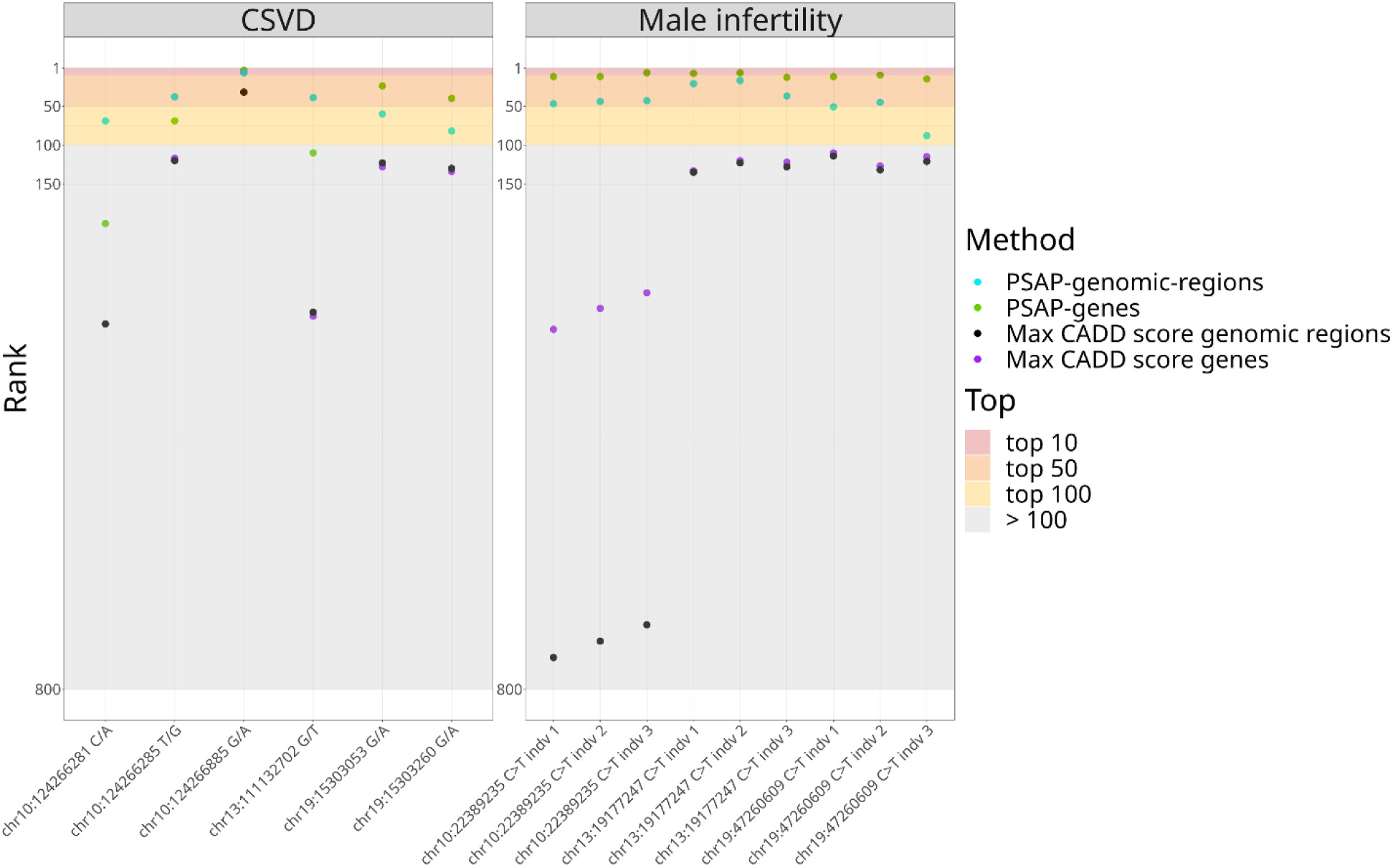
Prioritization of 6 known CSVD mutations and 3 male infertility candidate variants with PSAP-genomic-regions-CADD, PSAP-genes-CADD and the maximal CADD score on genes or CADD regions. The second dataset consisted of WGS data for 9 individuals from three families with clinically diagnosed male infertility (33). All causal variants fell within the top 20 of variants with prioritization by PSAP-genes-CADD, and within the top 50 for at least one case per family with PSAP-genomic-regions-CADD (within top 100 for all cases, Fig 4). PSAP-genomic-regions-CADD did not improve the ranking of these coding variants, which was expected considering the large number of variants in a WGS analysis (see S4 Table for the total number of variants in each analysis). The prioritization from PSAP-genomic-regions-CADD was still interesting to narrow the set of candidates for causal variants. In clinics when the CADD score alone is used, these variants would not have been prioritized (CADD score < 25, and rank > 100 with the maximal CADD score strategy). PSAP-genomic-regions-CADD thus allow a relevant prioritization of coding pathogenic variants in WGS sequencing and an unbiased exploratory analysis at the scale of the whole genome.

Using PSAP-genomic-regions-ACS or the ACS score alone, almost all of the CSVD and male infertility coding pathogenic variants had a rank greatly exceeding the top 100 (S4 Table). The only exception is one variant in *HTRA1* (10:124266885 G/A) that was ranked 3 by PSAP-genomic-regions-ACS and 10 by the maximal ACS score alone. This *HTRA1* variant was a splicing variant, which confirms the good performance of the PSAP-genomic-regions-ACS strategy on this type of variant.

## Discussion

Variant prioritization, especially in the case of very heterogeneous rare diseases, is a clinically-relevant methodological challenge for both clinicians and researchers. Mounting evidence suggests that current methods of analysis and their restriction to the coding genome are a hindrance to the discovery of new genetic variants implicated in rare diseases (16). We have developed PSAP-genomic-regions, an extension of the PSAP method to the whole genome using functionally-relevant genomic regions. PSAP-genomic-regions broadens the scope of variants evaluated by PSAP and addresses the issue of variant prioritization at an individual whole-genome scale.

PSAP-genomic-regions has been thoroughly tested and validated by using simulations emulating real-life scenarios of causal variant prioritization. PSAP-genomic-regions achieves a prioritization of coding pathogenic SNVs in the top 100 variants of an exome or genome which is a relevant number of variants to analyze for clinicians. Without use of prior knowledge on the disease, PSAP-genomic-regions achieves relevant variant prioritization within millions of variants to analyze, which is illustrated by the ranking of 6 variants involved in CSVD and 3 variants involved in familial cases of male infertility in the top 100 of WES and WGS data respectively. PSAP-genomic-regions thus helps with the diagnosis of such heterogeneous diseases in conjunction with other relevant information like the mode of transmission, prevalence or type of variant involved.

PSAP-genomic-regions also allows the scoring of variants otherwise discarded from the analysis, like splicing variants with a high predicted functional impact, and other non-coding variants of proven clinical significance. The only scenario for which PSAP-genomic-regions is not advantageous compared to the PSAP-genes strategy is for prioritizing coding variants in WGS data. In that case, using coding CADD regions, i.e. the coding parts of CADD regions for the analysis still yields better results compared to PSAP-genes (S7 Fig). Our simulations using known pathogenic variants have shown which PSAP strategy performs the best depending on the type of data and variant expected to be involved in the disease mechanism (S8 Fig). To effectively prioritize non-coding variants in WES and WGS, we advise the use of PSAP-genomic-regions-ACS. For coding variants, PSAP-genomic-regions-CADD gives the best results in WES, and PSAP-coding-genomic-regions-CADD performs best in WGS data. A two-step approach can also be carried out if there is no expected type of variant: first, the PSAP-genomic-regions-CADD or PSAP-coding-genomic-regions-CADD strategy is applied depending on the type of data, and if no coding variant of interest for the disease is found within the top results, PSAP-genomic-regions-ACS can be applied to look for non-coding variants of interest.

To the best of our knowledge, there is no other score of predicted pathogenicity for all possible SNVs comparable to CADD. Other methods have been developed to distinguish between coding pathogenic and neutral variants (34–39), but often restrict to non-synonymous variants. These methods were shown to perform better or have advantages compared to CADD for the limited set of variants they explore (34–39). Similar types of methods aim at prioritizing more constrained regions in the non-coding genome (18,20) or distinguishing deleterious non-coding variants from neutral ones (18,40). Other well-known methods for identification of pathogenic variants in exome and genome data rely on the use of HPO terms to make a prediction, like Exomiser (41) or Genomiser (30), making in comparison PSAP an unmatched prioritization tool. As any other bioinformatics variant prioritization method, it has to be used in conjunction with other lines of evidence to ultimately lead to any genetic diagnosis of a patient. PSAP-genomic-regions does not make assumption on the type of variants and does explore the whole genome. The ranking by p-values coming from the application of PSAP-genomic-regions to an individual’s variants is a useful way to narrow-down the list of variants to further investigate for both researchers and clinicians in different scenarios.

The method most comparable to the strategy followed by PSAP-genomic-regions is the recently-developed machine-learning algorithm FINSURF (42). FINSURF aims to predict the functional impact of non-coding variants in regulatory regions and has been applied to known pathogenic variants inserted in WGS data like we did. Nonetheless it has been difficult to compare properly the two methods considering FINSURF only scores non-coding variants in predefined regulatory regions, and the set of variants used to train the method is not available.

The main limitation of PSAP-genomic-regions comes from the score used to calibrate null distributions, namely the CADD score. We have observed that known pathogenic non-coding ClinVar variants that were not well-ranked by PSAP-genomic-regions had significantly lower CADD and ACS scores compared to splicing and better-ranked non-coding variants. Because such CADD score is likely to be seen in the general population, PSAP-genomic-regions will not be able to prioritize such a variant with at a low rank. We also observed that some CADD regions were badly-calibrated and resulted in the assignment of very low PSAP-genomic-regions p-values to putatively neutral variants in the 1000 Genomes Project. As allele frequencies from larger databases and more accurate pathogenicity scores become available, this will lead to an improvement of the PSAP method as well. The most recent release of the CADD score v1.7 (43) notably integrates regulatory annotations and may further improve the prioritization of non-coding pathogenic variants when integrated in PSAP-genomic-regions.

Many avenues of further development and improvement are open for PSAP-genomic-regions, including the inclusion and scoring of InDel variations and structural variants. Exploring the combination of the PSAP-genomic-regions p-values with other metrics or information coming from omics analysis could also improve prediction. Finally, the flexibility of the PSAP method makes it potentially adaptable to other more complex models like digenic and oligogenic models of inheritance, considering the increasing availability of information coming from gene networks and biological pathways.

## Materials and Methods

### Construction of PSAP null distributions

The first parameter is the units in which to construct the PSAP null distribution. Here we considered two unit strategies: the genes and the CADD regions (S1 Table). For the genes, the coding regions of genes were defined based on the biomaRt R package: the gene coding sequences were retrieved from Ensembl (44) by requesting the “genomic_coding_start” and “genomic_coding_end”, on both the hg19 and hg38 builds. To account for splicing regions, the coding regions were extended by two bases on both sides of the gene coding regions. In total, 19,780 genes were retrieved in hg19 and 23,163 in the hg38 build. For the CADD regions, their coordinates were downloaded from https://lysine.univ-brest.fr/RAVA-FIRST/ for the hg19 build and were lifted over to hg38 using the Ensembl Assembly Converter. CADD regions coordinates in hg38 are available on Easy-PSAP GitHub (https://github.com/msogloblinsky/Easy-PSAP).

There were 135,224 CADD regions in hg19 and 131,970 in hg38. For the coding CADD regions, i.e. the coding parts of CADD regions, we considered the intersection of the CADD regions and the gene coding regions for each build, which yielded 37,978 coding CADD regions in hg19 and 52,340 in hg38.

The second parameter is the allele frequencies database. Here we considered the global allele frequencies from the gnomAD database to calibrate the PSAP null distributions: gnomAD genome r2.0.1 for hg19 and gnomAD V3 (45) for hg38. For our purpose, we considered only single nucleotide variants (SNVs) annotated as PASS by the Variant Quality Score Recalibration (VQSR) of GATK (46) and located in well-covered regions. Well-covered regions in gnomAD genome were defined as regions for which 90% of individuals have coverage at depth 10. Variants not seen in gnomAD genome, not annotated as PASS or not located in well-covered regions (gnomAD genome version according to the build) have a frequency of 0 and thus did not contribute to the construction of the null distributions.

To ensure reliability of PSAP null distribution, it is crucial that the units are well covered in the database from which the allele frequencies are taken. Thus, we only considered units for which at least half of the unit was well-covered (as defined previously) in gnomAD genome (version according to the build). Coding regions of genes and well-covered regions in gnomAD genome were intersected to get the percentage of each gene’s coding regions that were well-covered in the database. The same steps were carried out with CADD regions as genomic units for PSAP, for hg19 and hg38 builds. PSAP null distributions were thus constructed for 19,283 and 18,395 genes in hg19 and hg38 respectively, 119,695 and 123,991 CADD regions, and 34,397 and 35,226 coding CADD regions in hg19 and hg38 respectively.

The third parameter is the pathogenicity score. Here, for the evaluation of PSAP on coding variants, we used the version 1.6 of CADD (47) for each build, accessible on the CADD website (https://cadd.gs.washington.edu/). For the evaluation on non-coding variants, which tend to have lower CADD scores than coding variants (48), we followed the strategy described in Bocher et al.(22) to adjust the RAW CADD score v1.6 of all possible SNVs on a PHRED scale stratifying by type of genomic regions: “coding”, “regulatory” and “intergenic”, resulting in “adjusted CADD scores”, referred to as “ACS”.

Easy-PSAP (https://github.com/msogloblinsky/Easy-PSAP) was used to generate null distributions according to the previously described input files and parameters. This resulted in 4 sets of null distributions for the AD and AR models for both hg19 and hg38 assemblies (S1 Table).

### Evaluating the performance of PSAP-genomic-regions using artificially-generated disease exomes and genomes

To evaluate the ability of PSAP-genomic-regions to prioritize known pathogenic variants in an individual, we leveraged artificially-generated disease exomes and genomes using available general population cohorts. These different PSAP strategies (see Table 1) were compared in terms of their performances to prioritize the known pathogenic variants.

The pathogenic ClinVar (25) SNVs with coordinates in hg19 and hg38 were downloaded from the NCBI website (https://www.ncbi.nlm.nih.gov/clinvar/, accessed on the 3rd of June 2022). Some of these ClinVar variants had an annotated mode of inheritance ("moi autosomal recessive" and "moi autosomal dominant"). From ClinVar, there were 12,776 variants annotated as AD and 12,776 variants annotated as AR. Variants were filtered out to keep only autosomal pathogenic SNVs having as review status either “reviewed by expert panel” or “criteria provided, multiple submitters, no conflicts”, which are the two best review status in ClinVar. There were 1,518 AD and 1,118 AR variants meeting these criteria.

For variants which did not have an annotated mode of inheritance, we used a curated version of the database OMIM, hOMIM (49) to retrieve a mode of inheritance, and kept variants that were always associated with an AD or AR mode of inheritance in hOMIM. The same filtering was applied, which left 3,641 additional variants for the AD and 1,706 for the AR model. In total, we had a set of 5,159 variants for the AD model and 2,824 variants for the AR model. Among these ClinVar variants, 4,965 and 2,680 variants were coding SNVs respectively for the AD and AR models. Similarly, 175 and 96 variants were non-coding variants for the AD model and AR models, among which 48 variants for the AD model and 64 for AR model fell within the boundaries covered by FREX exomes. The list of pathogenic ClinVar variants and their mode of inheritance can be found in S2 File.

We inserted each variant from our curated list of pathogenic ClinVar variants successively in each of the 533 high coverage genomes of Non-Finnish Europeans (NFE) from the 1000 Genomes Project phase 4 (NFE genomes) and each of the 574 exomes from the FREX project. An individual-focused QC was applied on both datasets using the RAVAQ R package (50): we performed a genotype and variant QC with default parameters corresponding to standard GATK hard filtering criteria, mean allele balance computed across heterozygous genotypes and call rates, except for MAX_AB_GENO_DEV = 0.25, MAX_ABHET_DEV, MIN_CALLRATE and MIN_FISHER_CALLRATE "disabled".

We conducted the artificially-generated disease genome and exome evaluation with PSAP null distributions in hg19 and hg38 respectively, to match with the build of the data. We then applied the 3 PSAP strategies mentioned previously (PSAP-genes-CADD, PSAP-genomic-regions-CADD and PSAP-genomic-regions-ACS). For each strategy, we kept the maximal pathogenicity score (CADD or ACS) for each unit (gene or CADD regions) and then ranked the units according to their PSAP p-value or to their pathogenicity score alone within each genome or exome. We compared the PSAP-genes-CADD and PSAP-genomic-regions-CADD strategies to using the maximal CADD score alone by gene or CADD regions, respectively; and the PSAP-genomic-regions-ACS strategy to using the maximal ACS score by CADD region. For each ClinVar variant, we retrieved its rank within each genome or exome. Coding ClinVar variants were evaluated with the 3 PSAP strategies whereas non-coding ClinVar variants were evaluated with the novel PSAP-genomic-regions-CADD and PSAP-genomic-regions-ACS strategies (see S2 Table for more details).

### Patient data analysis

The PSAP strategies were applied to real WES data from six unrelated patients affected by a CSVD for which the causal variant is known, which allowed a comparison of performance between the different strategies. The full description of the dataset can be found in [Aloui et al. 2021] (32), with the exception of the QC process. For this analysis, the same QC as for the FREX and 1000 Genomes Project datasets was performed. We applied PSAP-genes-CADD and PSAP-genomic-regions-CADD in hg19 to the six resolved CSVD patients’ exome data. The other PSAP parameters were the ones by default as described previously. Two of the individuals had a causal pathogenic variant in the gene *NOTCH3* (19:15303053 G/A and 19:15303260 G/A), one individual in the gene *COL4A2* (13:111132702 G/T) and three individuals in the gene *HTRA1* (10:124266285 T/G, 10:124266281 C/A and 10:124266885 G/A). The rank of the known CSVD variants among other heterozygote variants in the patient’s exome according to its PSAP p-value for the 2 strategies was then retrieved.

The PSAP strategies were also applied to WGS data of three families with clinically diagnosed forms of male infertility (33) and for which a pathogenic recessive variant was prioritized using a computational pipeline featuring the initial PSAP-genes implementation. Three affected individuals were analyzed for each family. The description of the whole dataset and candidate variant filtering process can be found in [Khan and Akbari et al. 2023] (33), except for the QC that was performed in the same way as for the CSVD data. Two other families were resolved from the same dataset, but considering that the causal variants were deletions we did not include them in the current analysis. The prioritized pathogenic variants were in the genes: *SPAG6* (chr10:22389235 C/T) for family 3, *TUBA3C* (chr13:19177247 C/T) for family 7 and *CCDC9* (chr19:47260609 C/T) for family 4. We applied PSAP–genes-CADD and PSAP-genomic-regions-CADD in hg38 to the 9 cases and retrieved the rank of the known male infertility variants among other homozygote variants in the patient’s genomes according to its PSAP p-value for the 2 strategies.

## Supporting Information captions

**S1 Fig. Summary statistics of pathogenicity scores and PSAP p-values (scale –log 10) for NFE individuals (one line by individual).**

**S2 Fig. Pathogenicity scores and PSAP p-values (scale –log 10) distributions for NFE individuals (maximal value for each genome), coding and non-coding ClinVar variants.**

**S3 Fig. Prioritization of splice variants versus other non-coding variants with PSAP on CADD regions with CADD or ACS**. P-values at 0 were replaced by a p-value of 10^-12^, which is lower than all the other non-zero p-values, for visualization purposes.

**S4 Fig. Distribution of CADD scores (A) and ACS scores (B) for ClinVar variants, by type of variant and mode of inheritance**. Coding: N=4,253 variants AD model and 2,245 variants AR model, Splicing: 102 variants AD model and 65 variants AR model, Non-coding top 100: 49 variants AD model and 19 variants AR model, Other non-coding: 24 variants AD model and 12 variants AR model.

**S5 Fig. Comparison of the the PSAP-genomic-regions strategy versus a pathogenicity score alone in artificially-simulated disease exomes**. Percentage of pathogenic non-coding and coding ClinVar variants reaching the top N of variants in at least 90% of FREX individuals, with PSAP-genomic-regions (darker shade of blue or green) or the pathogenicity score alone (lighter shade of blue or green), CADD or ACS (A) N = 48 non-coding AD variants and N = 64 non-coding AR variants (B) N = 4,965 coding AD variants and N = 2,680 coding AR variants.

**S6 Fig. Comparison of PSAP-genomic-regions-CADD and PSAP-genes-CADD for in artificially-simulated disease exomes.** Number of coding pathogenic ClinVar variants reaching the top N of variants in at least 90% of FREX individuals for each strategy.

**S7 Fig. Comparison of PSAP-coding-genomic-regions-CADD and PSAP-genes-CADD strategies for in artificially-simulated disease genomes.** Number of coding pathogenic ClinVar variants reaching the top N of variants in at least 90% of NFE individuals for each strategy.

**S8 Fig. Flowchart to choose the PSAP method of analysis depending on type of data and variants analyzed.**

**S1 Table. Currently available PSAP null distributions**. At https://lysine.univ-brest.fr/~msogloblinsky/share/data/).

**S2 Table. Strategies applied to construct and test PSAP null distributions.**

**S3 Table. Number and percentage of non-coding ClinVar variants in the top 10 of NFE genomes with PSAP-genomic-regions-CADD and PSAP-genomic-regions-ACS, by category of VEP consequence**. (A) Autosomal Dominant model (B) Autosomal Recessive Model.

**S4 Table. Ranks of 6 known CSVD variants and 3 male infertility candidate variants with PSAP-genes-CADD and PSAP-genomic-regions-CADD (1 row per individual).** Each CSVD variant was observed in a different individual. Each male infertility variant was observed in a different family consisting of three members each.

**S1 File. Method to generate PSAP null distributions.**

**S2 File. Pathogenic ClinVar variants used for the evaluation of PSAP null distributions through artificially-generated disease exomes and genomes.**

**S3 File. Rank of AD and AR non-coding non-splicing pathogenic ClinVar variants using PSAP-genomic-regions-ACS in artificially-generated disease genomes.**

**S4 File. Rank of Genomiser variants using PSAP-genomic-regions-ACS in artificially-generated disease genomes.**

## Notes

### Competing Interest Statement

The authors have declared no competing interest.

